# Null models for community dynamics: Beware of the cyclic shift algorithm

**DOI:** 10.1101/762278

**Authors:** Michael Kalyuzhny

## Abstract

**Aim:** Temporal patterns of community dynamics are drawing increasing interest due to their potential to shed light on assembly processes and anthropogenic effects. However, interpreting such patterns considerably benefits from comparing observed dynamics to the reference of a null model. For that aim, the cyclic shift permutations algorithm, which generates randomized null communities based on empirically observed time series, has recently been proposed. The use of this algorithm, which shifts each species time series randomly in time, has been justified by the claim that it preserves the temporal autocorrelation of single species. Hence it has been used to test the significance of various community patterns, in particular excessive compositional changes, biodiversity trends and community stability.

**Innovation:** Here we critically study the properties of the cyclic shift algorithm for the first time. We show that, unlike previously suggested, this algorithm does not preserve temporal autocorrelation due to the need to “wrap” the time series and assign the last observations to the first years. Moreover, this algorithm scrambles the initial state of the community, making any dynamics that results from deviations from equilibrium seem excessive. We exemplify that these two issues lead to a highly elevated type I error rate in tests for excessive compositional changes and richness trends.

**Conclusions:** Caution is needed when using the cyclic shift permutation algorithm and interpreting results obtained using it. Interpretation is further complicated because the algorithm removes all correlations between species. We suggest guidelines for using this method and discuss several possible alternative approaches. More research is needed on the best practices for using null models for temporal patterns.

## Introduction

One of the main approaches for the study of community ecology is documenting patterns of variation in ecological communities and underpinning the mechanistic basis for these patterns (Rosenzweig, 1995). Some of the most commonly studied patterns are species-area relationships (e.g. Preston, 1960; Rosindell & Cornell, 2007), latitudinal diversity gradients (e.g. Hillebrand, 2004; Usinowicz et al., 2017), productivity-diversity relationships (e.g. Kondoh, 2001; Tilman & Pacala, 1993) and so forth. Multiple explanations have been suggested for these patterns and studying them has shed light on the mechanisms determining species diversity in ecological communities (e.g. DeMalach, Zaady, & Kadmon, 2017; Usinowicz et al., 2017).

However, all the aforementioned patterns have a thing in common: they are static, representing a “snapshot” of ecological communities without any temporal dimension. While interest in these patterns continues, in recent years there is growing interest in understanding temporal patterns in communities (Dornelas et al., 2013; Loreau & de Mazancourt, 2013; Magurran, 2016; McGill, Dornelas, Gotelli, & Magurran, 2015). It is believed that some of these patterns, that represent community dynamics and assembly “in action”, may reveal new insights on the processes shaping ecological communities (Chisholm et al., 2014; Kalyuzhny, Seri, et al., 2014). For example, studies on the scaling of population fluctuations revealed the central role of temporal environmental variability in shaping ecological communities (Chisholm et al., 2014; Jabot & Lohier, 2016; Kalyuzhny, Schreiber, et al., 2014; Kalyuzhny, Kadmon, & Shnerb, 2015), and studies focusing on long-term changes in abundance and diversity revealed that population-level regulation is often weak (Kalyuzhny, Seri, et al., 2014; Knape & de Valpine, 2012; Ziebarth, Abbott, & Ives, 2010), while total-abundance and species diversity are indeed regulated (Brown, Ernest, Parody, & Haskell, 2001; Goheen, White, Ernest, & Brown, 2005; Gotelli et al., 2017; Magurran & Henderson, 2018). Moreover, in recent years there is an increasing interest in understanding richness trends and compositional turnover, partly motivated by concerns over the effect of anthropogenic activities on ecological communities (Elahi et al., 2015; Magurran et al., 2018; McGill et al., 2015; Vellend et al., 2013). Several studies have shown that while some local communities show richness trends, negative and positive changes may cancel out in multiple communities worldwide (Dornelas et al., 2014; Vellend et al., 2013). On the other hand, multiple ecological communities show large compositional turnover (Dornelas et al., 2014; Magurran et al., 2018).

This immediately raises the question: what qualifies as a “large” change in richness or composition? Stochastic community models generally predict that ecological communities would undergo constant changes at steady state (Lande, Engen, & Saether, 2003), and so do some deterministic models of nonlinear dynamics (May, 1976). This is true even for the simplest and most minimalistic models of community dynamics – Neutral Theory and Dynamic Equilibrium (DE) theory (Hubbell, 2001; MacArthur & Wilson, 1967). Consequently, and in analogy to null models of community patterns in space, temporal patterns should be compared to some null model to evaluate whether they deviate from the expectations under a minimalistic set of mechanisms (Gotelli & Graves, 1996; Gotelli & McCabe, 2002). Such null models preserve some aspects of the data and randomize others.

A promising suggestion for a null model of community dynamics is the cyclic shift permutations algorithm (Hallett et al., 2014), originally proposed for spatial analysis (Harms, Condit, Hubbell, & Foster, 2001). This algorithm gets as an input a matrix of species by years (or other temporal units). In each realization of the algorithm, the time series of every species is shifted forwards in time a random number of years *y*, independently of other species. The last *y* data points are then assigned to the first *y* years, hence “wrapping” the time series like a loop. For example, two possible resamples of the time series [1 2 3 4 5] could be [5 1 2 3 4] or [3 4 5 1 2] with equal probability. This approach has been claimed to preserve the autocorrelation structure and the abundance distribution of each species time series (Hallett et al., 2014; Lamy et al., 2019; Magurran et al., 2018), that result from the ecological dynamics of this species, while removing all correlations between species. Cyclic shift permutations have been used as a null model for richness and compositional changes (Demars et al., 2014; Magurran et al., 2018), changes in dominance (Jones & Magurran, 2018), and compensatory dynamics and stability of species diversity and total biomass (Gotelli et al., 2017; Hallett et al., 2014; Lamy et al., 2019; Magurran & Henderson, 2018). The application of the cyclic shift null model is greatly facilitated by the available implementation of this algorithm within the open source R package Codyn (Hallett et al., 2016).

Here we would like to point out two important issues with the use of cyclic shift permutations and investigate their implications for statistical tests of temporal patterns. We claim that a) Cyclic shift permutations do not preserve the autocorrelation structure of single species time series, especially the long-term autocorrelation patterns; and b) cyclic shift permutations scramble the initial state of the community, making any dynamics that result from initial deviations from equilibrium seem “excessive”. We show that these two properties lead to seriously inflated type I error rates when testing for excessive compositional changes and richness trends. For that aim, we generate synthetic community time series using two generic models, the independent single-species version of Dynamic Equilibrium theory (DE, MacArthur & Wilson, 1967; D. Simberloff, 1983; D. S. Simberloff, 1969) and Multispecies Ricker (Kalyuzhny & Shnerb, 2017; Kilpatrick & Ives, 2003). The former is a presence-absence dynamic model where species go extinct and arrive stochastically and independently of each other, while the latter considers abundances dynamics of interacting species in a fluctuating environment. For the synthetic time series, richness trends and compositional changes are compared to the predictions of cyclic shift permutations. We conclude by discussing the possible applications of the cyclic shift algorithm and other community null models.

### Do cyclic shift permutations preserve temporal autocorrelation?

The most important argument for using the cyclic shift permutations algorithm is that is (supposedly) preserves the temporal autocorrelation of the data. While this argument is highly intuitive, it ignores a crucial aspect of the algorithm – the “wrapping” procedure, where the last *y* data points (where *y* is the number of years that the time series has been shifted) are assigned to the first *y* years.

Consider the first and last year data points in the resampled time-series. It is highly likely that in the original time series, those were *consecutive* years, now maximally *separated* by the “wrapping” of the time series. Assuming the original time series had positive short-term temporal autocorrelation, this results in the last data point in the resample resembling considerably the first data point. This is generalizable, to some degree, to the first several data points resembling the final several data points. Moreover, the wrapping “attaches” a pair of years that were originally maximally separated in time, also distorting short-term autocorrelation.

Given data that was generated by some process, the goal of bootstrap resampling is generating more data that should resemble new data that would have been generated by repeating that process. Many ecological models, and DE and Ricker (for parameter regimes where the nonlinear effects do not take place) in particular, predict that consecutive time points would be relatively similar, and as time passes dissimilarity monotonously increases. Figure 1 exemplifies this for a single species undergoing stochastic colonization and extinction (Figure 1a), for multiple such species under DE (Figure 1b) and the Ricker model (Figure 1c). In all three cases, time series resampled using the cyclic shift permutations show a unimodal pattern of autocorrelation, very different from the original time series. Dissimilarity indeed initially increases, but then, after half the time series duration, begins to symmetrically decrease. Short term correlation is also quantitatively somewhat different from the original data.

**Figure 1.**
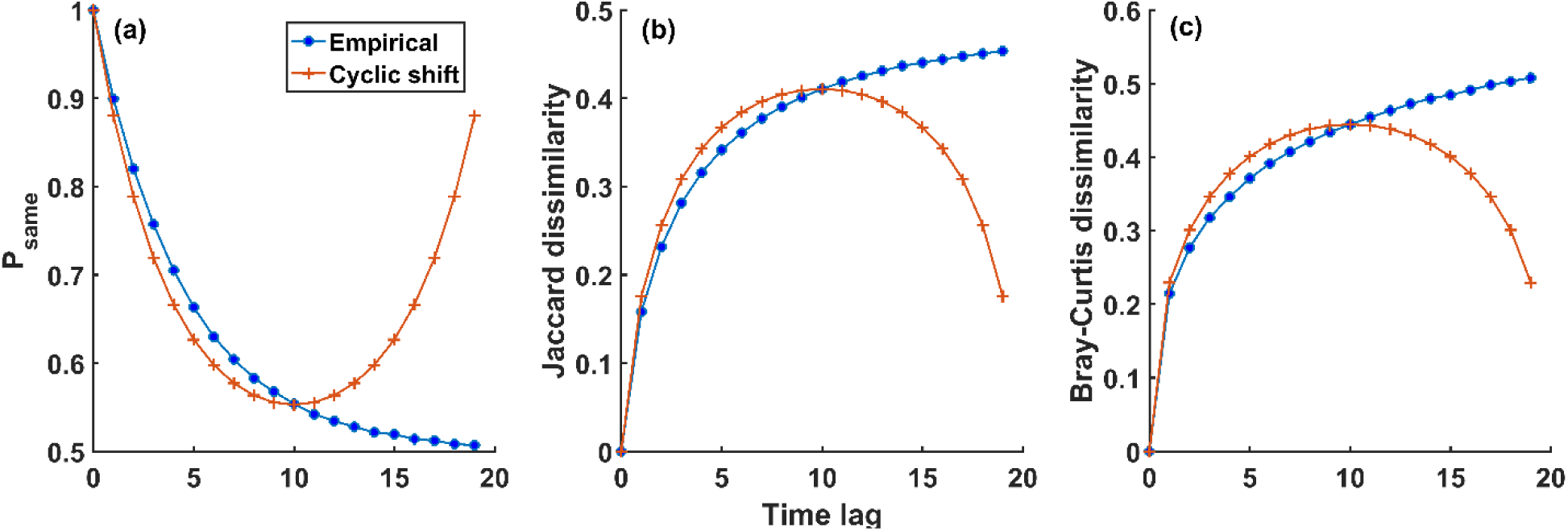
Comparison of temporal autocorrelation and dissimilarity patterns between “empirical” simulated time series and their resamples, obtained using the cyclic shift algorithm. In (**a**) 10^5^ colonization-extinction time series of a single species under Dynamic Equilibrium (DE) were simulated with colonization probability = extinction probability = 0.1. For each time series and time point, the probability to be at the same state (presence or absence) as the initial state was calculated for the “empirical” time series and for the resamples. In (**b**) and (**c**), 10^4^ communities of 100 and 10 species, respectively, were simulated under DE and multispecies Ricker. The Jaccard and Bray-Curtis compositional dissimilarity indices in every year were compared with the initial year for the empirical communities and resamples. For every empirical time series, 500 resamples were calculated and the results are averaged over the time series and resamples for every time point.

We conclude that the “wrapping” of the time series inherently distorts the temporal autocorrelation structure of ecological processes. This effect is dramatic on time scales that are on the order of the length of the time series, and less dramatic for shorter time scales. This raises the question – what are the implications of this, and of the “scrambling” of the initial state that was mentioned earlier, for the performance of statistical tests?

### Type I errors of richness trends and excessive turnover

We exemplified the consequences of these properties for the testing of two fundamental patterns of community dynamics – temporal trends in diversity and compositional turnover. These are quantified by calculating the linear regression slope of a) species richness and b) the dissimilarity of species composition of each year w.r.t the initial year; both versus time. These patterns have been studied in various communities and compared to the expectations under cyclic shift permutations to test for significance (Demars et al., 2014; Magurran et al., 2018). To examine the performance of the cyclic shift permutations null model in testing the significance of these patterns we generated 10^4^ synthetic communities under several parameter regimes using each of the DE and Ricker models. We then applied the cyclic shift algorithm 500 times to each community and compared the observed compositional and richness slopes in the time series generated by the model to the distribution of slopes under the cyclic shift algorithm. This allowed us to calculate the significance of the observed slopes for each community, and the proportion of significant (P < 0.05) results out of the generated communities.

Since simulations began at steady state, the percent of significant results (type I error) should be close to 0.05 if the null model is appropriate. This is particularly true for DE, since this model assumes that the species are independent, so the breaking of correlations imposed by cyclic shift permutations should have no effect. However, we found that for both models and under all parameter regimes, type I errors were considerably inflated (Figure 2).

**Figure 2.**
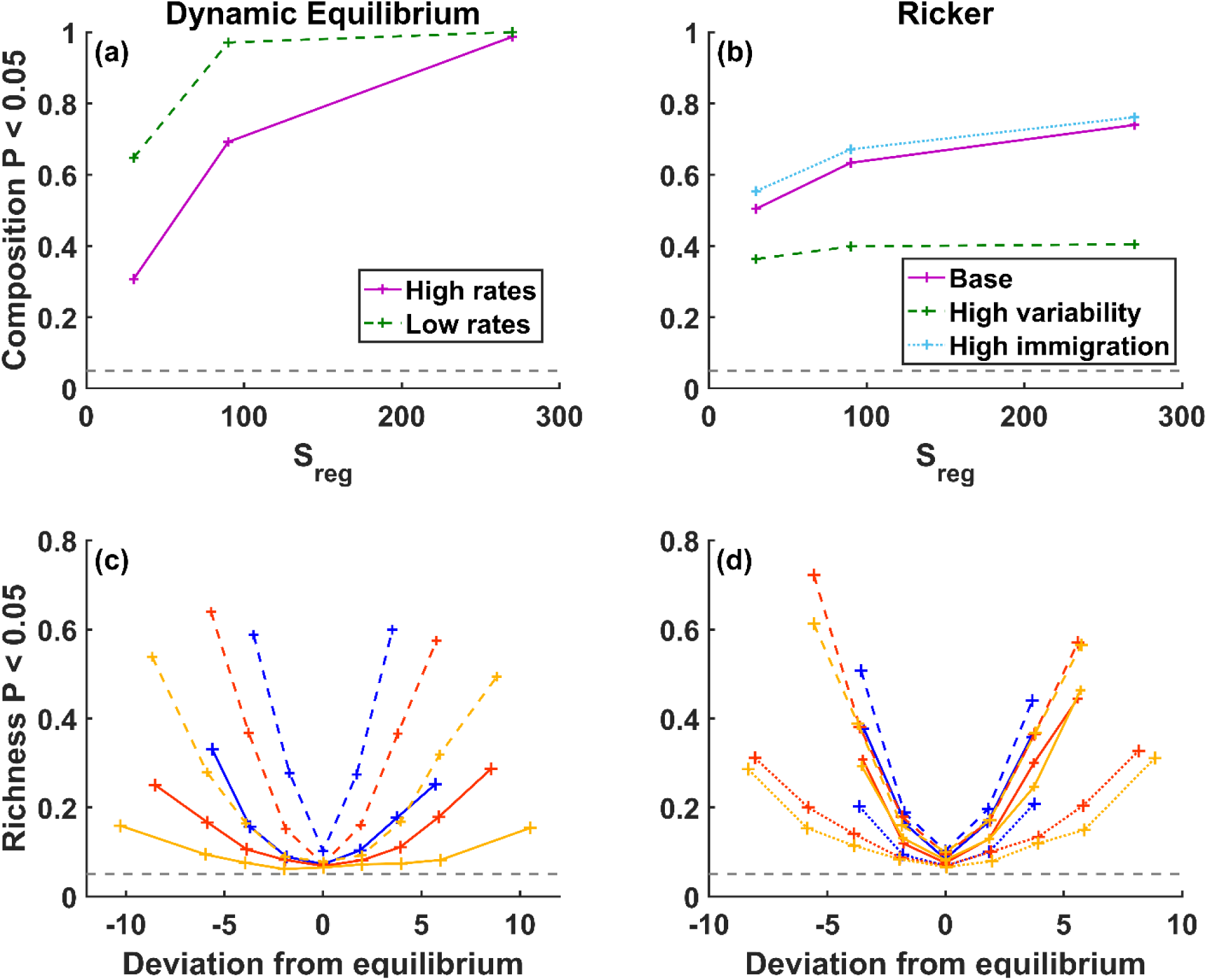
Proportion of significant tests (at α = 0.05) for excessive compositional changes (**a, b**) and richness trends (**c, d**) for data generated by the Dynamic Equilibrium (DE, **a, c**) and the multispecies Ricker (**b, d**) models. 10^4^ synthetic time series were generated for each parameter regime. For each time series, the slopes versus time of compositional dissimilarity w.r.t the first year and of specie richness were calculated. They were then compared with the distributions of slopes for 500 resamples (using the cyclic shift algorithm and a two sided test) of the synthetic time series to obtain a P value. In **c** and **d**, communities were assigned to bins based on the deviation of initial richness from equilibrium, and the proportion was calculated in each bin. We used the Jaccard and Bray-Curtis dissimilarity indices for DE and Ricker, respectively. The dashed grey line marks a proportion of 0.05.

Type I errors for compositional changes were very high (0.3 – 1) in all cases and tended to increase with the number of species in the pool (S_reg_, Figure 2a, b) and the number of species in the community. The latter is, at least in part, the reason that communities with more environmental fluctuations and less immigration, both with fewer species, having lower type I errors (Figure 2b). These results are the consequences of the unimodal pattern of dissimilarity with time that is generated by cyclic-shift permutations (Figure 1). The distribution of linear slopes fitted to this unimodal pattern is very different from the slope of the actual data, leading inevitably to a strong inflation of type I errors.

Regarding richness trends, communities starting at the equilibrium species richness had acceptable type I errors, with some exceptions (Figure 2c, d). For DE, communities with slow immigration and extinction rates and a regional pool of 30 species had a type I error probability of 0.102. For the Ricker model, communities with a high level of environmental fluctuations had type I error probabilities > 0.0945. However, the most pronounced result is that communities whose richness in the first year deviated from equilibrium, even by a few species, had a much higher type I error probability, reaching 0.3 – 0.7 in some parameter regimes (Figure 2c, d). Moreover, type I error probabilities increased sharply with the magnitude of the initial deviation from equilibrium. It is important to emphasize that we did not intentionally initiate the communities at a deviation from equilibrium richness. Rather, deviations were a result merely of stochastic dynamics at steady state. This implies that such deviations, and the resulting inflation in type I error, are to be expected in natural ecological communities.

The sharp increase in type I error rate as initial community richness moves away from equilibrium is a result of the “scrambling” of the initial state. Fairly generally, if some community property (richness, in this case, but without loss of generality) has an equilibrium value, and we find it initially at a different value, it is expected to return to that equilibrium. Hence, some level of short-term trends is to be expected under wide settings in ecological communities. However, the cyclic-shift permutation eliminates the initial state of the community, and as a result any trend would seem excessive. This effect is more pronounced if stochastic deviations are more likely and the rate of return to equilibrium is low, such as when colonization and extinction rates are low. This is also the reason for communities with low colonization and extinction rates having higher type I error rates (Figure 2a, c).

## Discussion

The cyclic shift algorithm is a generalistic and easy to apply null model, making it an appealing approach for testing a variety of different patterns. However, as we have shown, this approach has two fundamental undesired properties. First, it distorts the autocorrelation structure so that the end of the resampled time series for each species closely resembles its beginning. The short-term autocorrelation is also affected, but to a lesser degree. Moreover, the cyclic shift randomization “scrambles” the initial state of the community, making any dynamics that results from properties of this initial state seem unlikely. We have further exemplified that these fundamental limitations have severe consequences for type I error rates in tests for trends in species richness and excessive compositional changes. We believe that the fundamental nature of the issues with the cyclic shift null model would have negative consequences for tests that may be developed for other patterns as well.

Another aspect that must be considered carefully before using the cyclic shift algorithm is the implications of removing the correlations between species. If the goal of the analysis is testing for the significance of such correlations (e.g. Hallett et al., 2014) then using a null with no correlations definitely makes sense. However, other aspects of the dynamics, such as temporal changes in diversity, composition and dominance, may very well be affected by correlations between species. These correlations could be caused by biotic interactions or responses to environmental changes, and the interpretation of finding excessive changes compared to such a null of independent species should be carefully considered. Even more so, one should be cautious about using cyclic-shift permutations on data that is available at a resolution of less than a year (e.g. Magurran et al., 2018; Magurran & Henderson, 2018). In such data, correlation between species may be the result of seasonality, and randomly shifting each species independently of other species removes its effect. Consequently, the likely strong effects of seasonality on community composition will be detected as excessive changes.

These issues do not lessen, however, the need that led to the development of the cyclic shift algorithm. Indeed, we believe that the interest in temporal patterns will continue to grow, along with the need for a null model to serve as reference for them. We would like to suggest several possible directions for addressing this need.

First, it is possible that the performance of the cyclic shift algorithm as a null model is not as bad for some patterns. Because of the concerns we raise here, we believe that it is up to the future researchers who wish to use this approach for testing some pattern to convince that its performance is reasonable. We recommend specifically examining its performance on the statistical test of interest applied to simulated data, as we did. One should ensure that the characteristics of the simulated data, such as the rates of the dynamics and total richness, resemble the empirical patterns.

An alternative approach that has indeed been suggested as a null model for community dynamics is neutral models (Dornelas et al., 2014; Gotelli & McGill, 2006; Hubbell, 2001). However, it has been shown that neutral models where stochastic events affect individuals independently (known as “demographic stochasticity” or “ecological drift”), such as the classical Unified Neutral Theory of Biodiversity and Biogeography (Hubbell, 2001), predict considerably smaller changes then observed in multiple communities (Dornelas et al., 2014; Kalyuzhny, Schreiber, et al., 2014; Kalyuzhny, Seri, et al., 2014). It has been shown that this is the result of ignoring environmental fluctuations, which affect the growth rate of entire populations synchronously (Chisholm et al., 2014; Fung, O’Dwyer, Rahman, Fletcher, & Chisholm, 2016; Jabot & Lohier, 2016; Kalyuzhny et al., 2015). Hence, a neutral model with environmental fluctuations would be a much more appropriate null (Jabot & Lohier, 2016; Kalyuzhny et al., 2015). Furthermore, neutral theories impose compensatory dynamics, or negative correlations, between species, which stem from the zero-sum assumption. This makes them not necessarily the best choice as a null model (Gotelli & McGill, 2006). Another alternative would be to try to fit multispecies autoregressive models (Ives, Dennis, Cottingham, & Carpenter, 2003). This framework is more flexible, and one may decide to preserve (or not to preserve) multiple properties such as the autocorrelation structure of the data, the magnitude of fluctuations, the correlations between species and the initial state of the community. This approach has not been studied much as a null model to this day.

Finally, a null model for presence-absence data named Presence-Absence Resampling wIthin periodS (PARIS) has recently been suggested as a methodology to generate synthetic communities where each species independently undergoes colonization and extinction dynamics at fixed rates (Kalyuzhny, Flather, Shnerb & Kadmon, 2019). While this approach imposes the autocorrelation structure of a Poisson process and species independence, it preserves the initial state of the community and has recently been shown to have excellent statistical properties for data satisfying these assumptions (Kalyuzhny, Flather, Shnerb & Kadmon, 2019). PARIS is also very easy to apply to ecological time series because, like the cyclic-shift algorithm, it is a randomization-based methodology.

Overall, we believe that studying temporal patterns has great promise to shed light on the processes shaping ecological communities. This promise is amplified by the increasing availability of extensive datasets of temporal community dynamics (Dornelas et sl., 2018). However, more research is required on how to appropriately analyze such data, and in particular on the best practices for applying null models of community dynamics. We hope this work will reduce the improper application of such null models and help guide the use and development of more appropriate null models for temporal community dynamics.

## Methods

### Models

We have studies the statistical performance of the cyclic shift permutations algorithm by applying it to synthetic time series generated using two models: the independent-species version of Dynamic Equilibrium theory (DE, D. Simberloff, 1983; D. S. Simberloff, 1969) and a multispecies Ricker model (M. Kalyuzhny & Shnerb, 2017; Kilpatrick & Ives, 2003).

DE is the simplest possible model of community dynamics, considering a local community that receives immigrants from a regional pool with *S*_*reg*_ species. Time is modeled in discrete time steps. Every time step, if species *i* is present, it has a fixed probability *e*_*i*_ to go extinct and be absent by the next time step, while if it is absent it will arrive and be present by the next time step with probability *c*_*i*_. The probability of any species to be present at steady state is 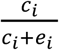, hence communities are initialized at steady state by drawing the presence of each species from a Bernoulli distribution with the aforementioned probability.

Despite its simplicity, this model has two parameters (*c*_*i*_ and *e*_*i*_) for each species. For generating them, we assumed that colonization and extinction rates follow a lognormal distribution with both mean and variance *M*. This is a reasonable assumption since both body mass and species abundance are often approximately lognormally distributed (Brown, Marquet, & Taper, 1993; Ulrich, Ollik, & Ugland, 2010). The rates are drawn independently and transformed to probabilities by the 1 - exp(-*k*) transformation, where *k* is a rate. This transformation calculates the probability for the occurrence of an event in a Poisson process. For figure 1 we took M = 0.2 and for figure 2 we considered two regimes, one with high and one with low rates, which are M = 0.4 (median probability of event: 0.21) and 0.08 (median probability of event: 0.022), respectively.

The second model we used is a stochastic, discrete-time multispecies Ricker model. In this model, the expected population of species *i* at time *t*+1, *N*_*i,t*+1_, in the absence of immigration is:

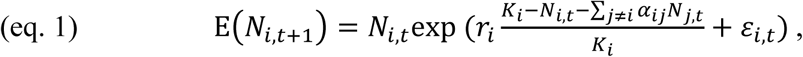

where *r*_*i*_ and *K*_*i*_ are the growth rate and carrying capacity of species i, respectively, *α*_*ij*_ is the per capita effect of an individual of species j on the growth of species i, representing inter-specific interactions, and *ε*_*i,t*_ represents stochastic fluctuations in the growth rate due to environmental changes. *ε*_*i,t*_ is normally distributed with a mean of 0 and variance of *σ*_*e*_^2^.

While eq. 1 represents the expected population of species i at time t, the actual population size is drawn from a Poisson distribution: *N*_*i,t*+1_∼Poisson(E(*N*_*i,t*+1_)). This introduces demographic stochasticity, that is, random variation between individuals in demography, as well as the discreteness of individual, which allows species to go stochastically extinct. Finally, after the local demography step described above, we introduce a Poisson distributed number of immigrants (with mean *I*, representing immigration rate), which are chosen uniformly from the *S*_*reg*_ species available in the species pool.

For generating the parameters of the model, we assumed that the *K*_*i*_s of species are lognormally distributed with mean and SD of 1000, the *r*_*i*_s are exponentially distributed with mean 0.5 and the *α*_*ij*_s are gamma distributed with mean and SD of 0.1. We considered three parameter regimes. In the basal regime, σ_e_^2^ = 0.1 and I = 2. In the high temporal variability regime, we changed σ_e_^2^ to 0.5 and in the high immigration regime we changed I to be 10. Simulations were initialized with each species at its *K*_*i*_ and run for 500 time steps to equilibrate.

For all models, we considered three levels of S_reg_: 30, 90 and 270 species and the duration of the time series was 20 time steps (after equilibration).

### Statistical tests

We are interested in testing the performance of tests for excessive compositional turnover and richness trends. For compositional turnover, we computed the Jaccard dissimilarity (for DE) or Bray-Curtis dissimilarity (for Ricker) of every year with respect to the initial year (as in Figure 1b, c) and used the linear slope of dissimilarity vs. time as the test statistic. For richness trends, we computed the slope of the regression of richness vs. time. For every synthetic community generated by DE or Ricker we calculated these two test statistics, generated 500 resampled communities by applying cyclic shift permutations to the original data, and then compared the observed statistics to their distribution in the resampled communities using a two-sided test. This gave us the P value of both statistics for every community. This procedure is in line with the approach of Magurran et al. (2018).

To evaluate the performance of the statistics, we calculated the proportion of significant results (using α = 0.05). Since we expect no excessive changes, this proportion can be thought of as type I error rate, which should not exceed α. For examining the tests of compositional change (Figure 2a, b), the proportion was calculated over all 10^4^ synthetic communities in a given parameter regime. For examining the tests of richness trends, we first assigned the communities to bins according to their initial deviation from equilibrium richness. Equilibrium richness was calculated as the average richness in the 20 year data for the sake of simplicity and resemblance to empirical analyses, where the real equilibrium is unknown. The bins that were used were D ≤ -7, -7 < D ≤ -5, -5 < D ≤ -3, -3 < D ≤ -1, -1 < D ≤ 1, 1 < D ≤ 3, 3 < D ≤ 5, 5 < D ≤ 7, 7 < D, where *D* is initial deviation from richness equilibrium. In each bin, we calculated the average D (presented on the X axis), and the proportion of significant results among the communities in the bin (presented on the Y axis). Bins with less than 30 communities were discarded.

All analyses were performed in Matlab 2016a with the full code supplied in supporting information S1.

## Acknowledgements

I thank Ronen Kadmon and Nadav Shnerb for encouraging discussions of the results. M.K. is supported by the Adams Fellowship Program of the Israel Academy of Sciences and Humanities.

