## supporting information S1 for "Null models for community dynamics: Beware of the cyclic shift algorithm": analysis_fig1.m

```
%generate single-species colonization-extinction dynamics

ex1 = .1; %extinction rates
im1 = .1; %immigration rates
len1 = 20;

ittr = 100000;
cyclic_res_ittr = 500;

%compute 'ittr' realizations of the dynamics:
comp_r1_l1 = MWDR_2_mex(im1*ones(ittr,1) , ex1*ones(ittr,1) , binornd(ones(ittr,1) , im1/(im1 + ex1)) , 1 , len1);

%compute the probability of change for that dynamics:
pchange_r1_l1 = zeros(1,len1);
for tt = 1:len1
    pchange_r1_l1(tt) = sum(comp_r1_l1(tt,:) == comp_r1_l1(1,:)) / ittr;
end


%resample using cyclic shift and compute average 
pchange_r1_l1_res = zeros(cyclic_res_ittr,len1);
parfor rr = 1:cyclic_res_ittr;
    
    comp_cyclic = cyclic_shift(comp_r1_l1);
    
    for tt = 1:len1
        pchange_r1_l1_res(rr,tt) = sum(comp_cyclic(tt,:) == comp_cyclic(1,:)) / ittr;
    end
    
end

save('fig1_results.mat')

%% generate community time series for DE:

%input parameters:
sreg = 100;
rate_mu = .4; %avarage rates
rate_sigma = .4; %variance (NOT STD!) of rates
ittr = 10000;
cyclic_res_ittr = 500;
len1 = 20;

%transform the parameters of the lognormal into parameters of the asociated
%normal
mx1_e=log((rate_mu^2)/sqrt(rate_sigma+rate_mu^2));
vx1_e=sqrt(log(rate_sigma/(rate_mu^2)+1));

%output:
J_emps = zeros(ittr,len1);
J_cyclic_res = zeros(ittr,len1);

parfor ii = 1:ittr
    

    %generate rates and transfor into probabilities:
    im_rates = lognrnd(mx1_e,vx1_e,1,sreg);
    ex_rates = lognrnd(mx1_e,vx1_e,1,sreg);
    im = 1 - exp(-im_rates)'; %transform into probabilities
    ex = 1 - exp(-ex_rates)';
    
    %generate data:
    dyn = MWDR_2_mex(im , ex , binornd(ones(sreg,1) , im./(im + ex)) , 1 , len1);
    J_emps(ii,:) = J_init(dyn);
    
    %resample:
    res_now = zeros(cyclic_res_ittr,len1);
    
    for rr = 1:cyclic_res_ittr
        dyn_res = cyclic_shift(dyn);
        res_now(rr,:) = J_init(dyn_res);
    end
    %average resamples:
    
    J_cyclic_res(ii,:) = mean(res_now);
end

save('fig1_results.mat')

%% generate community time series for Ricker:

%input parameters:
r_mean = 0.3;
K_mean = 1000;
K_var = 1000;
A_mean = 0.1;
A_std = 0.1;
env_sigma = 0.5;
ims = 1;
S_reg = 30;
t_equi = 500;
t_obs = 20;
ittr = 10000;
cyclic_res_ittr = 500;

mx1_e=log((K_mean^2)/sqrt(K_var+K_mean^2));
vx1_e=sqrt(log(K_var/(K_mean^2)+1));

%output:
BC_emps = zeros(ittr,t_obs);
BC_cyclic_res = zeros(ittr,t_obs);
   
parfor ii = 1:ittr
    
    rs = exprnd(r_mean,S_reg,1);
    Ks = lognrnd(mx1_e,vx1_e,S_reg,1);
    A = gamrnd((A_mean/A_std)^2,(A_std^2)/A_mean,S_reg,S_reg);
    N0 = Ks;
    
    N = multi_ricker3_3_mex(rs,Ks,A,env_sigma,ims,N0,t_equi+t_obs);
    N = N';
    N = N(t_equi+1:end,:);
    
    BC_emps(ii,:) = BC_init(N);
    
    %resample:
    res_now = zeros(cyclic_res_ittr,t_obs);
    
    for rr = 1:cyclic_res_ittr
        dyn_res = cyclic_shift(N);
        res_now(rr,:) = BC_init(dyn_res);
    end
    
    %average resamples:
    BC_cyclic_res(ii,:) = mean(res_now);

end

save('fig1_results.mat')

%% generate plot:

figure()

s1 = subplot(1,3,1);
hold on
set(s1,'FontSize',18,'FontWeight','bold','LineWidth',1.5);
plot(0:len1-1 , pchange_r1_l1 , '-o' ,'MarkerFaceColor',[0 0 1],'LineWidth',1.5)
plot(0:len1-1 , mean(pchange_r1_l1_res) , '-+', 'MarkerSize' ,8 ,'LineWidth',1.5 )
ylabel('P_s_a_m_e')
legend({'Empirical' , 'Cyclic shift'})
text(1,.98, '(a)' ,'FontSize',18,'FontWeight','bold')

s1 = subplot(1,3,2);
hold on
set(s1,'FontSize',18,'FontWeight','bold','LineWidth',1.5);
plot(0:len1-1 , mean(J_emps) , '-o' ,'MarkerFaceColor',[0 0 1],'LineWidth',1.5)
plot(0:len1-1 , mean(J_cyclic_res) , '-+', 'MarkerSize' ,8 ,'LineWidth',1.5 )
ylabel('Jaccard dissimilarity')
xlabel('Time lag')
text(1,.48, '(b)' ,'FontSize',18,'FontWeight','bold')


s1 = subplot(1,3,3);
hold on
set(s1,'FontSize',18,'FontWeight','bold','LineWidth',1.5);
plot(0:len1-1 , mean(BC_emps) , '-o' ,'MarkerFaceColor',[0 0 1],'LineWidth',1.5)
plot(0:len1-1 , mean(BC_cyclic_res) , '-+', 'MarkerSize' ,8 ,'LineWidth',1.5 )
ylabel('Bray-Curtis dissimilarity')
text(1 , .58, '(c)' , 'FontSize' , 18, 'FontWeight' , 'bold')
```
