## supporting information S1 for "Null models for community dynamics: Beware of the cyclic shift algorithm": analysis_fig2.m

```
%% prepare the simulations of DE:

%the results are put into the arrays DER (Dynamic Equilibrium Results) which
%has three dimensions: (rates(2),Sreg(3),itterations)

tic

S_reg_levels = [30 90 270];
rates_levels = [.4, .08];
ittr = 10000;
cyclic_res_ittr = 500;
len1 = 20;

DER_alpha_p = nan(length(rates_levels),length(S_reg_levels),ittr); %p values
DER_beta_p = nan(length(rates_levels),length(S_reg_levels),ittr); %p values
DER_sr1 = nan(length(rates_levels),length(S_reg_levels),ittr); %initial richness
DER_srm = nan(length(rates_levels),length(S_reg_levels),ittr); %mean richness

for rr = 1:length(rates_levels)
    
    %transform the parameters of the lognormal into parameters of the asociated normal
    mx1_e=log((rates_levels(rr)^2)/sqrt(rates_levels(rr) + rates_levels(rr) ^ 2));
    vx1_e=sqrt(log(rates_levels(rr)/(rates_levels(rr)^2)+1));
    
    for ss = 1:length(S_reg_levels)
        for ii = 1:ittr
            [rr ss ii]
            %generate probabilities:
            im_rates = lognrnd(mx1_e,vx1_e,1,S_reg_levels(ss));
            ex_rates = lognrnd(mx1_e,vx1_e,1,S_reg_levels(ss));
            im = 1 - exp(-im_rates)'; %transform into probabilities
            ex = 1 - exp(-ex_rates)';

            %generate data:
            dyn = MWDR_2_mex(im , ex , binornd(ones(S_reg_levels(ss),1) , im./(im + ex)) , 1 , len1);
            sr_real = sum(dyn,2);
            
            %calculate changes in alpha and beta:
            alpha_trend = polyfit(1:len1,sr_real',1);
            alpha_trend = alpha_trend(1);
            J_emp = J_init(dyn);
            beta_trend = polyfit(1:len1,J_emp,1);
            beta_trend = beta_trend(1);
            
            %resample:
            alpha_trend_res = nan(1,cyclic_res_ittr);
            beta_trend_res = nan(1,cyclic_res_ittr);
            
            parfor jj = 1:cyclic_res_ittr
                %generate data:
                dyn_res = cyclic_shift(dyn);
                sr_null = sum(dyn_res,2);
                
                %calculate resamples stats:
                tmp = polyfit(1:len1,sr_null',1);
                alpha_trend_res(jj) = tmp(1);
                J_res = J_init(dyn_res);
                tmp = polyfit(1:len1,J_res,1);
                beta_trend_res(jj) = tmp(1);
                
            end
            
            %output:
            DER_alpha_p(rr,ss,ii) = dist_pval(alpha_trend_res,alpha_trend);
            DER_sr1(rr,ss,ii) = sr_real(1);
            DER_srm(rr,ss,ii) = mean(sr_real);
            DER_beta_p(rr,ss,ii) = dist_pval(beta_trend_res,beta_trend);
        end
    end
end

toc

%% generate plots of delta_s vs p:
%calculate the dependence of proportions of rejection on deviation from
%initial richness

min_gr = [-1000 -7 -5 -3 -1 1 3 5 7 1000];
thresh_group = 30; %this is the minimal number of observations in a group to be considered. any group with less observations will be ignored.

DER_delta = DER_sr1 - DER_srm; %the deviations from the mean

obs_gr_DE = zeros(length(rates_levels),length(S_reg_levels),ittr); %which group does each observation belong to?
cat_obs_DE = zeros(length(rates_levels) , length(S_reg_levels) , length(min_gr)-1); %how many observations in each category?
cat_avg_DE = nan(length(rates_levels) , length(S_reg_levels) , length(min_gr)-1); %avg delta (deviation form the average) in each group
cat_sig_DE = nan(length(rates_levels) , length(S_reg_levels) , length(min_gr)-1); %proportion significant in each group
        
for rr = 1:length(rates_levels)
    for sss = 1:length(S_reg_levels)
        
        for ii = 1:ittr
            obs_gr_DE(rr,sss,ii) = find(DER_delta(rr,sss,ii) <= min_gr, 1) - 1;
        end

        %for the figure of delta_s vs p, for each categoty compute number of
        %observations, avg. richness and proportion significant:

        [cat_obs_DE(rr,sss,:),~] = hist(reshape(obs_gr_DE(rr,sss,:),1,ittr) , 1:length(min_gr)-1); %number of obs. in each category.

        for cc = 1:length(min_gr)-1
            obs_this_group = reshape(obs_gr_DE(rr,sss,:),1,ittr) == cc;
            
            %average richness in categories:
            res = reshape(DER_delta(rr,sss,:),1,ittr);
            cat_avg_DE(rr,sss,cc) = mean(res(obs_this_group));

            %percent significant in categories:
            res = reshape(DER_alpha_p(rr,sss,:),1,ittr);
            cat_sig_DE(rr,sss,cc) = sum(res(obs_this_group) < 0.05)/cat_obs_DE(rr,sss,cc);

        end
    end
end

cat_sig_DE(cat_obs_DE < thresh_group) = NaN; %remove groups with too few observations

save('fig2_DE_results.mat')

%% simulations of the Ricker Model:

%the results are put into the arrays RR (Ricker Results) which
%have three dimensions: (regime(3),Sreg(3),itterations). there are three
%regimes: 1. baseline. 2. hight env. variability. 3. high immigration
tic

r_mean = 0.5;
K_mean = 1000;
K_var = 1000^2;
A_mean = 0.1;
A_std = 0.1;
env_sigma_levels = [0.1 0.5];
ims_levels = [2 10];
S_reg_levels = [30 90 270];
t_equi = 500;
ittr = 10000;
cyclic_res_ittr = 500;
len1 = 20;

%transform the parameters of the lognormal into parameters of the asociated normal
mx1_e=log((K_mean^2)/sqrt(K_var+K_mean^2));
vx1_e=sqrt(log(K_var/(K_mean^2)+1));

RR_alpha_p = nan(3,length(S_reg_levels),ittr); %p values
RR_beta_p = nan(3,length(S_reg_levels),ittr); %p values
RR_sr1 = nan(3,length(S_reg_levels),ittr); %initial richness
RR_srm = nan(3,length(S_reg_levels),ittr); %mean richness

for rr = 1:3
    
    %what parameter regime are we in?
    if rr == 1
        ims = ims_levels(1);
        env_sigma = env_sigma_levels(1);
    elseif rr ==2
        ims = ims_levels(1);
        env_sigma = env_sigma_levels(2);
    elseif rr == 3
        ims = ims_levels(2);
        env_sigma = env_sigma_levels(1);
    end

    for ss = 1:length(S_reg_levels)
        for ii = 1:ittr
            [rr ss ii]
            
            %generate community properties:
            S_reg = S_reg_levels(ss);
            rs = exprnd(r_mean,S_reg,1);
            Ks = lognrnd(mx1_e,vx1_e,S_reg,1);
            A = gamrnd((A_mean/A_std)^2,(A_std^2)/A_mean,S_reg,S_reg);
            N0 = Ks;

            %generate data:
            N = multi_ricker3_3_mex(rs,Ks,A,env_sigma,ims,N0,t_equi + len1);
            N = N';
            N = N(t_equi+1:end,:);
            sr_real = sum(N > 0,2);
            
            %calculate changes in alpha and beta:
            alpha_trend = polyfit(1:len1,sr_real',1);
            alpha_trend = alpha_trend(1);
            BC_emp = BC_init(N);
            beta_trend = polyfit(1:len1,BC_emp,1);
            beta_trend = beta_trend(1);
            
            %resample:
            alpha_trend_res = nan(1,cyclic_res_ittr);
            beta_trend_res = nan(1,cyclic_res_ittr);
            
            parfor jj = 1:cyclic_res_ittr
                %generate data:
                dyn_res = cyclic_shift(N);
                sr_null = sum(dyn_res > 0 , 2);
                
                %calculate resamples stats:
                tmp = polyfit(1:len1,sr_null',1);
                alpha_trend_res(jj) = tmp(1);
                BC_res = BC_init(dyn_res);
                tmp = polyfit(1:len1,BC_res,1);
                beta_trend_res(jj) = tmp(1);
                
            end
            
            %output:
            RR_alpha_p(rr,ss,ii) = dist_pval(alpha_trend_res,alpha_trend);
            RR_sr1(rr,ss,ii) = sr_real(1);
            RR_srm(rr,ss,ii) = mean(sr_real);
            RR_beta_p(rr,ss,ii) = dist_pval(beta_trend_res,beta_trend);
        end
    end
end

toc

%% diveide into categories:

min_gr = [-1000 -7 -5 -3 -1 1 3 5 7 1000];
thresh_group = 50; %this is the minimal number of observations in a group to be considered. any group with less observations will be ignored.

RR_delta = RR_sr1 - RR_srm; %the deviations from the mean

obs_gr_R = zeros(3 , length(S_reg_levels),ittr); %which group does each observation belong to?
cat_obs_R = zeros(3 , length(S_reg_levels) , length(min_gr)-1); %how many observations in each category?
cat_avg_R = nan(3 , length(S_reg_levels) , length(min_gr)-1); %avg delta (deviation form the average richness) in each group
cat_sig_R = nan(3 , length(S_reg_levels) , length(min_gr)-1); %proportion significant in each group
        
for rr = 1:3
    for sss = 1:length(S_reg_levels)
        
        for ii = 1:ittr
            obs_gr_R(rr,sss,ii) = find(RR_delta(rr,sss,ii) <= min_gr, 1) - 1;
        end

        %for the figure of delta_s vs p, for each categoty compute number of
        %observations, avg. richness and proportion significant:

        [cat_obs_R(rr,sss,:),~] = hist(reshape(obs_gr_R(rr,sss,:),1,ittr) , 1:length(min_gr)-1); %number of obs. in each category.

        for cc = 1:length(min_gr)-1
            obs_this_group = reshape(obs_gr_R(rr,sss,:),1,ittr) == cc;
            
            %average richness in categories:
            res = reshape(RR_delta(rr,sss,:),1,ittr);
            cat_avg_R(rr,sss,cc) = mean(res(obs_this_group));

            %percent significant in categories:
            res = reshape(RR_alpha_p(rr,sss,:),1,ittr);
            cat_sig_R(rr,sss,cc) = sum(res(obs_this_group) < 0.05)/cat_obs_R(rr,sss,cc);

        end
    end
end

cat_sig_R(cat_obs_R < thresh_group) = NaN; %remove groups with too few observations

save('fig2_R_results.mat')

%% generate figure:

cols_sreg = [0 0 1; 1 0.2 0; 1 0.7 0]; %the colors for the different levels of S_reg used in the two lower panels
col_regime = [.75 0 .75; 0 .5 0; .3 .75 .93 ];

figure()
s1 = subplot(2,2,1);
hold(s1,'on')
colormap(col_regime);

set(s1,'FontSize',18,'FontWeight','bold','LineWidth',1.5 ,'ColorOrder',col_regime);
su = sum(DER_beta_p < 0.05,3)/ittr;
plot(S_reg_levels , su(1,:),'-+','LineWidth',1.5);
plot(S_reg_levels , su(2,:),'--+','LineWidth',1.5);
ylabel('Composition P < 0.05')
xlabel('S_r_e_g')
title('Dynamic Equilibrium')
legend({'High rates', 'Low rates'})
ylim([0 1])
xlim([0 300])
line([0 300],[0.05 0.05],'LineStyle','--','Color',[.5 .5 .5],'LineWidth',1.5)
text(5 , .95, '(a)' , 'FontSize' , 18, 'FontWeight' , 'bold')

s1 = subplot(2,2,2);
hold(s1,'on')
set(s1,'FontSize',18,'FontWeight','bold','LineWidth',1.5,'ColorOrder',col_regime);
su = (sum(RR_beta_p < 0.05,3)/ittr);
plot(S_reg_levels , su(1,:) ,'-+','LineWidth',1.5);
plot(S_reg_levels , su(2,:) ,'--+','LineWidth',1.5);
plot(S_reg_levels , su(3,:) ,':+','LineWidth',1.5);


xlabel('S_r_e_g')
title('Ricker')
legend({'Base', 'High variability' 'High immigration'})
ylim([0 1])
xlim([0 300])
line([0 300],[0.05 0.05],'LineStyle','--','Color',[.5 .5 .5],'LineWidth',1.5)
text(5 , .95, '(b)' , 'FontSize' , 18, 'FontWeight' , 'bold')


s1 = subplot(2,2,3);
hold(s1,'on')
set(s1,'FontSize',18,'FontWeight','bold','LineWidth',1.5);
for sss = 1:length(S_reg_levels)
    plot(reshape(cat_avg_DE(1,sss,:),1,length(min_gr)-1) , reshape(cat_sig_DE(1,sss,:),1,length(min_gr)-1), '-+' , 'MarkerEdgeColor', cols_sreg(sss,:) , 'Color' , cols_sreg(sss,:) , 'LineWidth',1.5, 'MarkerSize' ,8)
    plot(reshape(cat_avg_DE(2,sss,:),1,length(min_gr)-1) , reshape(cat_sig_DE(2,sss,:),1,length(min_gr)-1), '--+' , 'MarkerEdgeColor', cols_sreg(sss,:) , 'Color' , cols_sreg(sss,:) , 'LineWidth',1.5)
end
ylabel('Richness P < 0.05')
xlabel('Deviation from equilibrium')
xlim([-12 12])
ylim([0 0.8])
line([-12 12],[0.05 0.05],'LineStyle','--','Color',[.5 .5 .5],'LineWidth',1.5)
text(-11.6 , .76, '(c)' , 'FontSize' , 18, 'FontWeight' , 'bold')


s1 = subplot(2,2,4);
hold(s1,'on')
set(s1,'FontSize',18,'FontWeight','bold','LineWidth',1.5);
for sss = 1:length(S_reg_levels)
    plot(reshape(cat_avg_R(1,sss,:),1,length(min_gr)-1) , reshape(cat_sig_R(1,sss,:),1,length(min_gr)-1), '-+' , 'MarkerEdgeColor', cols_sreg(sss,:) , 'Color' , cols_sreg(sss,:) , 'LineWidth',1.5)
    plot(reshape(cat_avg_R(2,sss,:),1,length(min_gr)-1) , reshape(cat_sig_R(2,sss,:),1,length(min_gr)-1), '--+' , 'MarkerEdgeColor', cols_sreg(sss,:) , 'Color' , cols_sreg(sss,:) , 'LineWidth' , 1.5, 'MarkerSize' ,8)
    plot(reshape(cat_avg_R(3,sss,:),1,length(min_gr)-1) , reshape(cat_sig_R(3,sss,:),1,length(min_gr)-1), ':+' , 'MarkerEdgeColor', cols_sreg(sss,:) , 'Color' , cols_sreg(sss,:) , 'LineWidth',1.5 , 'MarkerSize' ,8)
end
xlabel('Deviation from equilibrium')
xlim([-10 10])
ylim([0 0.8])
line([-10 10],[0.05 0.05],'LineStyle','--','Color',[.5 .5 .5],'LineWidth',1.5)
text(-9.7 , .76, '(d)' , 'FontSize' , 18, 'FontWeight' , 'bold')
```
