## supporting information S1 for "Null models for community dynamics: Beware of the cyclic shift algorithm": BC_init.m

```
function [BCT] = BC_init(composition)
%BC_init calculates the Bray-Cuttis dissimilarity of every time step compared to the first.
%Compisition is a 2D array of abundances where each row is a year and each
%column is a species.

t_tot=length(composition(:,1));
BCT=zeros(1,t_tot);

for t=1:t_tot
    BCT(t) = sum(abs(composition(t,:) - composition(1,:))) / sum(composition(t,:) + composition(1,:));
end

end
```
