## supporting information S1 for "Null models for community dynamics: Beware of the cyclic shift algorithm": cyclic_shift.m

```
function [dyn_res] = cyclic_shift(dyn)
%in the input, 'dyn', a row is a year and a column is a species

[years, sps] = size(dyn);

dyn_res = nan(years,sps);

for sp = 1:sps
    shift = unidrnd(years)-1; %how many places to the right to shift the first year?
    inds = (1:years)';
    inds_shifted = inds + years - shift;
    too_big = inds_shifted > years;
    inds_shifted(too_big) = inds_shifted(too_big) - years;
    dyn_res(:,sp) = dyn(inds_shifted,sp);
end

end
```
