## supporting information S1 for "Null models for community dynamics: Beware of the cyclic shift algorithm": multi_ricker3_3.m

```
function [N]=multi_ricker3_3(r,K,A,env_sigma,ims,N0,t_tot)

%assuming uniform regional SAD and ims<1

% This function simulates the Multispecies Ricker process. All vectors are
% column vectors, env_sigma is the STD of the environmental noise, N0
% represents initial condition and t_tot is the total timesteps to run.
% 'ims' represents the average of immigrants added each year. The number
% iteslf is drawn from a poisson distribution, and the species pool is
% assumed to be uniform. The function outputs both absolute population size
% (N).

%since we are using relative interactions, turn the diagonal of the
%interactions matrix to ones

for i=1:length(N0)
    A(i,i)=1;
end


sps=length(N0); %total number of species
N=zeros(sps,t_tot);
P=zeros(sps,t_tot);
N(:,1)=N0;
P(:,1)=N(:,1)/sum(N(:,1));

for t=2:t_tot
    N(:,t)=poissrnd2((N(:,t-1).*exp(r.*(ones(sps,1)-(A*N(:,t-1)./K))+normrnd(0,env_sigma,sps,1)))); %intrinsic dynamics
    add_ims = poissrnd(ims);
    for ii = 1:add_ims;
        arrival=randi(sps,1);
        N(arrival,t)=N(arrival,t)+1; 
    end
end

end
```
