## supporting information S1 for "Null models for community dynamics: Beware of the cyclic shift algorithm": poissrnd2.m

```
function [res]=poissrnd2(param)
%generates a vector of length m of poisson random numbers using the normal
%approximation. receives a vector of length 'm' of parameters
len=length(param);
res=zeros(1,len);

to_approx=find(param>50);
not_to_approx=find(param<=50);
res(to_approx)=round(normrnd(param(to_approx),sqrt(param(to_approx))));
res(not_to_approx)=poissrnd(param(not_to_approx));
end
```
