## supporting information S1 for "Null models for community dynamics: Beware of the cyclic shift algorithm": README.docx

### How to use this code?

This code is written in Matlab (2016).

The two master scripts that perform all analyses are ‘analysis_fig1.m’ and ‘analysis_fig2.m’, which call all other functions in turn. Parallel computation, as well as Matlab Coder – the ‘multi_ricker3_3’ is run in the code with the suffix _mex to accelerate the code.
